# Critical evaluation of kinetic schemes for coagulation

**DOI:** 10.1101/2021.04.12.439461

**Authors:** Alexandre Ranc, Salome Bru, Simon Mendez, Muriel Giansily-Blaizot, Franck Nicoud, Rodrigo Méndez Rojano

**Affiliations:** Department of Haematology Biology, CHU, Univ Montpellier, Montpellier, France; IMAG, Univ Montpellier, CNRS, Montpellier, France; Polytech, Univ Montpellier, Montpellier, France

**Keywords:** Modeling of the coagulation cascade, Hemostasis, Contact system, TF-pathway

## Abstract

Computational models of the coagulation cascade are used for a wide range of applications in bio-medical engineering such as drug and bio-medical device developments. However, a lack of robustness of numerical models has been highlighted when studying clinically relevant scenarios. In order to develop more robust models, numerical simulations need to be confronted with realistic situations relevant to clinical practice. In this work, two well-established numerical representations of the coagulation cascade initiated by the intrinsic and extrinsic systems, respectively, were compared with thrombin generation assays considering realistic pathological conditions. Proper modifications were needed to align the *in vitro* and *in silico* data, namely; adapting initial conditions to the thrombin assay system, omitting reactions irrelevant to our case study, and improving the fitting of some reaction rates. The modified models were able to capture the experimental trends of thrombin generation for a range of concentrations of factors XII, XI, and VIII for cases in which the coagulation cascade is triggered through the extrinsic and intrinsic systems. Our work emphasizes that when existing coagulation cascade models are extrapolated to experimental settings for which they were not calibrated, careful adjustments must be made. We show that the two coagulation models used in this work can predict physiological conditions, but when studying pathological conditions, proper modifications are needed to improve the numerical results.

## 1 Introduction

In recent decades, numerical modeling of the coagulation cascade has been used to understand the complex dynamics of thrombin formation [1–3]. These models have allowed the discovery of coagulation factor interactions [4], to study the pharmacokinetics of new drugs [5], and investigate the dynamics of thrombus formation in pathological conditions such as aortic aneurysms or infarcted left ventricles [6,7]. More recently, a computational thrombosis model was used to understand the underlying dynamics that control bleeding severity among hemophilia A patients highlighting the ability of numerical models to construct realistic synthetic data sets to overcome the absence of data in clinical practice [8].

Numerical representations of the coagulation cascade aim to mimic the thrombin generation process which is the result of the balance between prothrombin conversion and thrombin inactivation [9], thrombin being the key enzyme of the blood clotting cascade. Thrombin generation can be summarized *in vivo* with the following enzymatic steps based on [10]. The tissue factor (TF) is released by damaged vessels or prothrombotic cells including tumor cells, activated mono-nuclear cells, or micro-vesicles. It complexes with the activated factor (F) VII (FVIIa) and activates the FX either directly or indirectly through the FIX. Then, the activated FX activates the prothrombin into the key thrombin factor. Prothrombin conversion is self-regulated. Thrombin activates the FXI and both the co-factors VIII and V of factors IX and X, respectively, creating feedback loops that increase its generation, leading to the thrombin burst. In parallel, thrombin cleaves the fibrinogen into fibrin monomers forming an unstable network which is subsequently stabilized by activated FXIII. On the other hand, the coagulation cascade can also be triggered by the contact activation system as a result of the interactions of FXII, kininogen, and prekallikrein with negatively charged surfaces. This coagulation route, also known as the intrinsic pathway, was historically neglected due to the absence of clinical phenotype in cases of FXII deficiency but has now a renewed interest due to promising anti-FXI and anti-FXII drugs to limit thrombosis in situations where blood comes into contact with artificial surfaces of medical devices [11].

In order to represent the aforementioned cascade of reactions, computational models contain a large number of input parameters that have uncertainties related to their experimental characterization and a lack of standardization in the community when inferring the reaction rate values [12]. Therefore, it is not surprising to find reaction rate values spanning several orders of magnitude for the same coagulation reaction [13]. This lack of robustness in computational models is illustrated when mathematical models are used to study pathological conditions. As pointed out by Chelle *et al.* [14], most of the existing computational models fail to reproduce the thrombin formation dynamics in clinical settings without prior parameter optimization. To avoid this issue, some authors have suggested that instead of trying to find a universal set of reactions, reduced coagulation models are enough to reproduce the coagulation dynamics [9,15,16]. Nevertheless, a prior calibration step is still needed. A strategy to increase the robustness of coagulation models can be to assess the sensitivity of thrombin formation to coagulation factor deficiencies. By doing such sensitivity studies, the robustness of the reaction scheme can be tested more deeply, and the reaction rate values can be tuned in a more relevant way, making sure that the final model is not only able to represent one regime, but rather a whole variety of situations. In addition, predicting thrombin kinetics under pathological conditions (like factor deficiencies) is the ultimate goal of most mathematical models.

In the present work, we evaluated two well-established mathematical models of the coagulation cascade from Chatterjee *et al.* [2] for the contact pathway and from Hockin *et al.* [1] and Butenas *et al.* [3] for the TF pathway. Three clotting factor deficiencies were studied to investigate if the models could reproduce the dynamics of thrombin generation. For each model, the numerical simulations were compared with experimental data using calibrated automated thrombography (CAT) [17]. A range of decreasing concentrations of the coagulation factor FVIII was considered in order to represent the hemophilia A condition. In addition, deficient FXII and FXI plasmas were used to probe the initiation of the coagulation cascade with the intrinsic pathway, which is of special interest to medical devices. The comparison of numerical and experimental data allowed us to quantify the accuracy of the numerical models, identify limitations, and suggest adjustments that improved the predictive capabilities of the kinetic schemes.

Section 2 describes the methods for experimental thrombography and numerical simulations. In Section 3.1 experimental and numerical results for a range of concentrations of FXII, FXI, and FVIII triggered through the intrinsic pathway are presented. Section 3.2 shows the results for the range of FVIII concentrations triggered with TF pathway. Finally, Section 4 discusses the results and the limitations of our work.

## 2 Materials and Methods

### 2.1 Plasma Samples

Samples used to characterize the contact pathway were derived from lyophilized plasmas. To avoid inter-individual physiological variations of clotting factor levels, lyophilized Standard Human Plasma (SHP^®^) [Siemens Healthcare, Erlangen, France] constituted of industrial pooled citrated platelet poor plasmas (PPP) and obtained from at least one hundred healthy donors was used. Lyophilized factor immunodepleted plasmas were industrial pooled citrated PPP with a qualified specific factor activity lower than 1% [Siemens Healthcare, Erlangen, Germany]. According to manufacturer specifications, donors were pretested for PT, APTT, factor II, V, VII, VIII, IX, X, XI, XII levels, which were all within the normal range. SHP was diluted with lyophilized factor immunodepleted plasmas (FXII, FXI, and FVIII), in order to obtain a range of final concentrations from less than 1% to 100% (100%, 50%, 15%, 5%, 1% and < 1% treated as 0% from here on). Silica, mixed with rabbit cephalin in STA^®^-PTT-A^®^ (Stago, Asnires-sur-Seine, France), was used to initiate the intrinsic pathway.

Inherited deficient frozen plasmas were preferred to lyophilized plasmas to assess the TF coagulation pathway because of higher values for the thrombin generation assay (TGA) parameters (data not shown). Duchemin *et al.* [18] highlighted the same difference between the lyophilized and immunodepleted haemophiliac plasmas. Frozen plasmas used to assess the TF coagulation pathway were derived from a single hemophilia A patient (Cryopep, Montpellier, France). To obtain a range of FVIII concentrations from less than 1% to 100% (100%, 50%, 15%, 5%, 1% and < 1%), the hemophilia A plasma was picked with increased amounts of in-house fresh frozen PPP plasmas from seven healthy donors tested for normal PT, APTT and fibrinogen levels. To avoid any residual platelets, two sequential centrifugations were performed at 2250 g at 20 °C for 10 minutes. The inhouse PPP aliquots were stored at −80 °C. The TF coagulation pathway was triggered using the PPP-Reagent LOW (Thrombinoscope^®^ BV, Maastricht, The Netherlands) containing a final TF concentration of 1pM. Low concentrations of TF were chosen to obtain a thrombin generation mainly dependent upon feedback activation loops and FVIII and FIX clotting factors [19].

### 2.2 Calibrated automated measurement of thrombin generation

Thrombin generation was performed in PPP using Fluoroscan Ascent^®^ (FluCa kit, Thrombinoscope^®^, Synapse BV, Maastrich, The Netherlands), according to the method described by Hemker *et al.* [17] that measures the thrombin generated thanks to a fluorochrome. 80 *μ*L of plasma were mixed with 20 *μ*L of STA^®^-PTT-A^®^ or PPP reagent LOW, according to the coagulation pathway tested, and 20 *μ*L of fluorescent reagent FluCa kit. The latter contains the calcium chloride which is mandatory to activate the coagulation cascade and the Fluo-Substrate which is a fluorogenic substrate mixed with the FluoBuffer. Fluorescence intensity was detected at wavelengths of 390 nm (excitation filter) and 460 nm (emission filter), every 20 seconds. Each individual sample was analyzed in triplicate simultaneously with a thrombin calibrator as reference for a stable thrombin activity of approximately 600 nM. The calibrator enables the conversion of the fluorescence signal into thrombin concentration. The signal was treated to correct inner filtering effects, such as substrate consumption and abnormal plasma color. Analyses were conducted on Immulon 2HB round-bottom 96-well plates (Stago - Asnieres-sur-Seine, France).

The parameters calculated by Thrombinoscope^®^ software included: (i) endogenous thrombin potential (ETP) to evaluate the overall effect on thrombin generation, (ii) the IIa_*max*_ which corresponds to the largest value of thrombin, (iii) the time to peak (*τ_max_*) which is the time required to reach IIa_*max*_ [17]. The physical interpretation of these quantities is provided in Fig. 1.

**Fig. 1.**
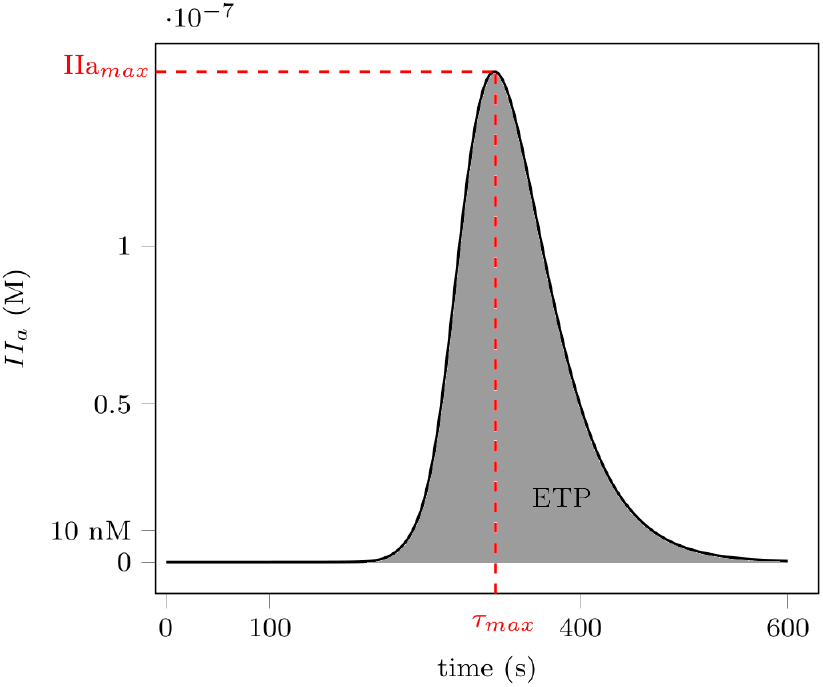
Variables used for *in vitro* and *in silico* comparison: IIa_*max*_, *τ_max_* and ETP for a thrombin formation curve in time.

### 2.3 Kinetic Schemes and Mathematical modeling

Two coagulation kinetic models were used in the current study. For the sake of simplicity, the models of Chatterjee *et al.* [2] and Butenas *et al.* [3] are named Int and Ext, respectively, making reference to the intrinsic and extrinsic pathway.

The models were implemented in the in-house YALES2BIO^1^ software. YALES2BIO is a mathematical tool dedicated to study blood flows. This software has recently been used to study the kinetics of thrombin formation in medical devices, triggered by the contact activation system [20,21]. To solve each biochemical scheme, a set of differential equations was obtained by applying the law of mass action to the biochemical reactions belonging to each kinetic model. This representation mimics a closed homogeneous reactor in which the only mechanisms that influence the dynamics of coagulation factors are production and consumption (without diffusive or convective transport of species [22]). The system of equations was then computationally solved to obtain the evolution of coagulation factors in time using the initial conditions displayed in Table 1. Initial conditions were computed from the dilutions used in TGA assays. The nominal initial conditions used in the study were derived accounting for the dilution steps of the calibrated automated thrombinography.

**Table 1.**
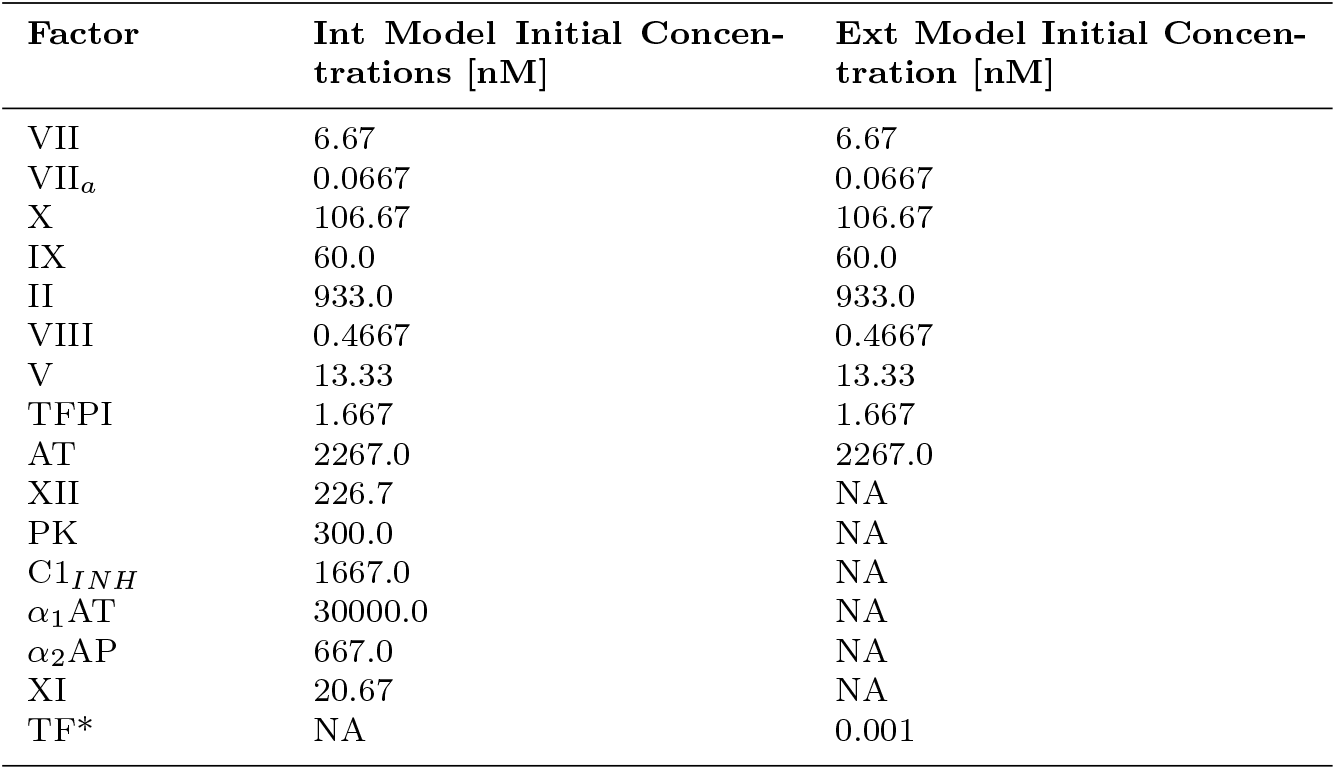
Initial conditions of Int and Ext models, derived from TGA dilution. * TF nominal value, see Section 3.2.

**Table 2.**
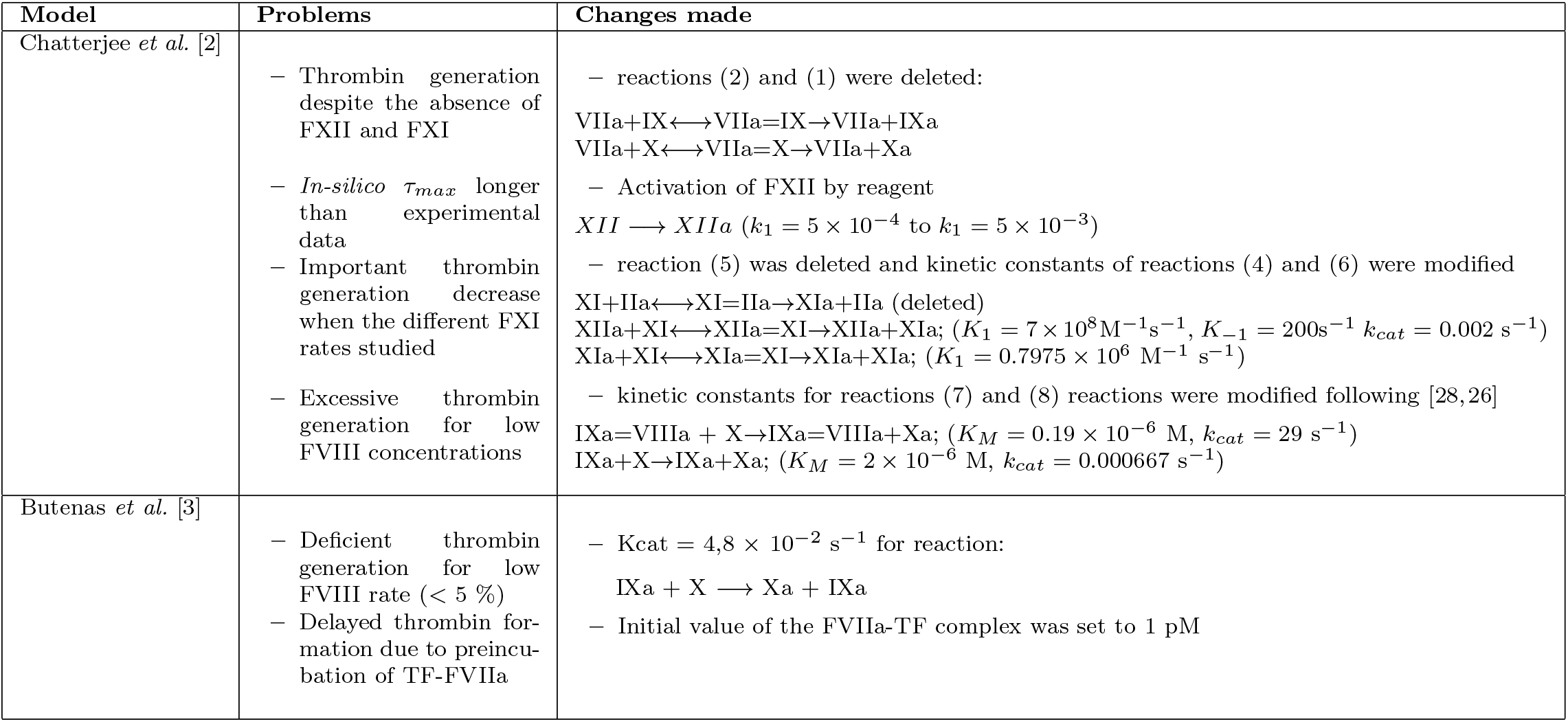
Modifications performed for Int and Ext models.

The Int model was published in 2010 and describes the coagulation triggered by the contact pathway [2]. It was built upon the existing model of Hockin *et al.* [1] with additions concerning the contact pathway reactions. The model is triggered by contact activation of factor XII. Since factor immunodepleted plasmas were collected without corn trypsin inhibitor (CTI), the reaction 35 that describes the inhibition of factor XIIa by CTI in Chatter-jee *et al.* [2] was suppressed. Moreover, since the thrombin generation is the basis of the analysis performed in the present study, fibrin-related reactions (# 50 to 57 from [2]) were deleted to save computing time without interfering with the thrombin generation. The Ext model was described in the literature by Hockin *et al.* [1] and upgraded in 2004 by Butenas *et al.* [3] with a supplementary reaction that describes the activation of factor X by the activated factor IX (IXa).

The Int and Ext models were modified to align some reactions and kinetic schemes to the studied physiological conditions. The modified formulations were compared with the original numerical results and experimental data from TGA. The rationale for each specific modification is discussed in Section 3. Both the original and modified kinetic schemes used are listed in the Supplementary Information.

### 2.4 *In vitro* vs in *silico* comparison

To establish a comprehensive comparison between the *in silico* and *in vitro* results, the ETP, IIa_*max*_, *τ_max_* variables were considered (see Fig. 1). Relative values were computed considering the physiological production of thrombin in PPP as a reference. In the case of Fig. 3 and Fig. 7, these values are used in order to evaluate the general trend for the different concentrations rather than comparisons using the absolute values.

## 3 Results

### 3.1 Int model for the contact pathway

TGA experiments and simulations using the original Int model were conducted over a range of decreasing concentrations of FXII, FXI, and FVIII. Figure 2 shows the numerical results and experimental TGA data for plasma with 100%, 15%, and 0% for FXII, FXI and FVIII. Four main discrepancies are observed:

– Numerical data at a concentration of 100% show a delay and steeper rise in thrombin formation compared to experimental data.
– When the concentration is less than 100%, the original model cannot capture the trend observed experimentally.
– In the numerical cases with 0% of FXII the coagulation cascade should not start at all, but the model predicts thrombin generation.
– For the low concentration of FVIII (0 %) numerical thrombin production is significantly larger than experimental data.

**Fig. 2.**
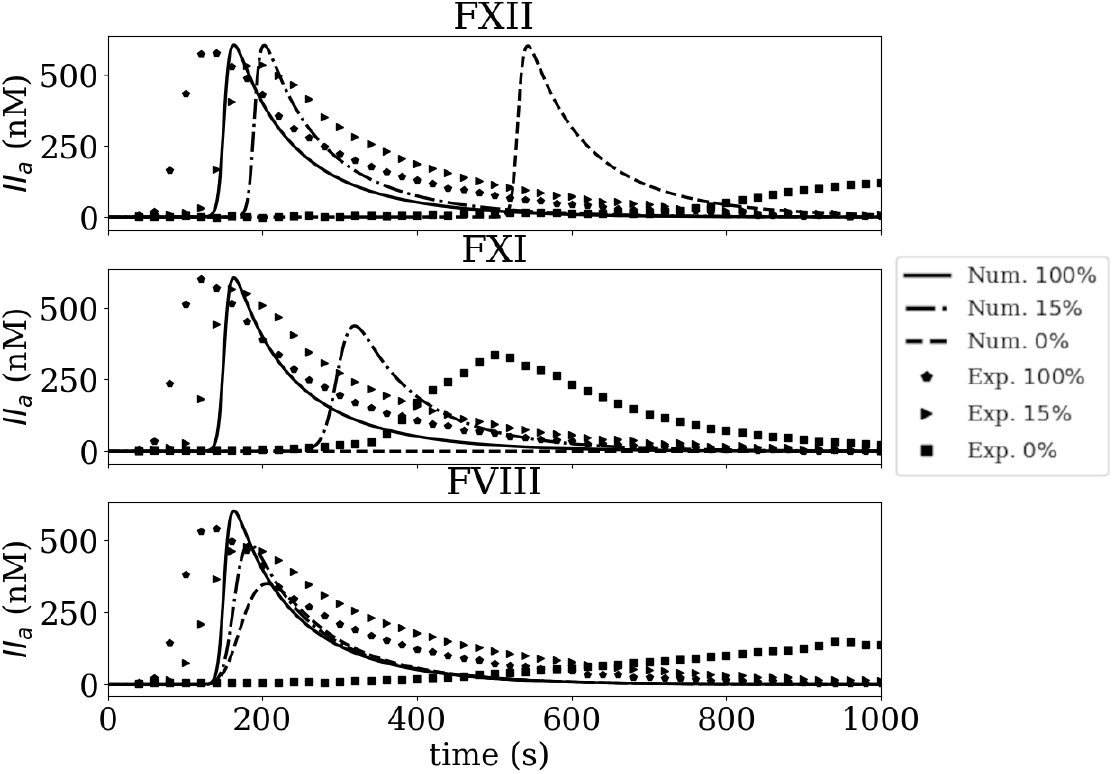
Experimental thrombin formation (TGA) and numerical thrombin formation using the original Int model for FXII, FXI and FVIII at 100%, 15%, and 0%.

Figure 3 shows the evolution of relative ETP, IIa_*max*_, *τ_max_* values for all the FXI, FXII, and FVIII concentrations: 0, 1, 5, 15, 50, and 100%. The original and modified numerical model results are shown along with the experimental data. It can be observed that for the FXI case both ETP and IIa*_max_* values predicted by the original model show a descending trend steeper than the experimental data. In terms of *τ_max_,* the values computed by the original model are significantly larger than the experimental data. Looking at FXII, the *τ_max_* trend is better captured by the original model, however, ETP and IIa_*max*_ values do not follow the same trend observed experimentally for low concentrations of FXII. In addition, undesired thrombin production is observed for FXII 0% concentration. It is worth saying that *in vitro* thrombin formation for FXII and FXI at 0% cases can be explained by the presence of residual amounts of coagulation factors in the respective immunodepleted plasmas, however, this should not happen numerically. The ETP and IIa_*max*_ values computed by the original model for FVIII concentrations follow a similar trend as the experimental data, nonetheless, the increased *τ_max_*. values observed experimentally for decreasing concentrations of factor VIII are not observed at the same level in the numerical data for the original model. The modifications made to improve the model are discussed below.

**Fig. 3.**
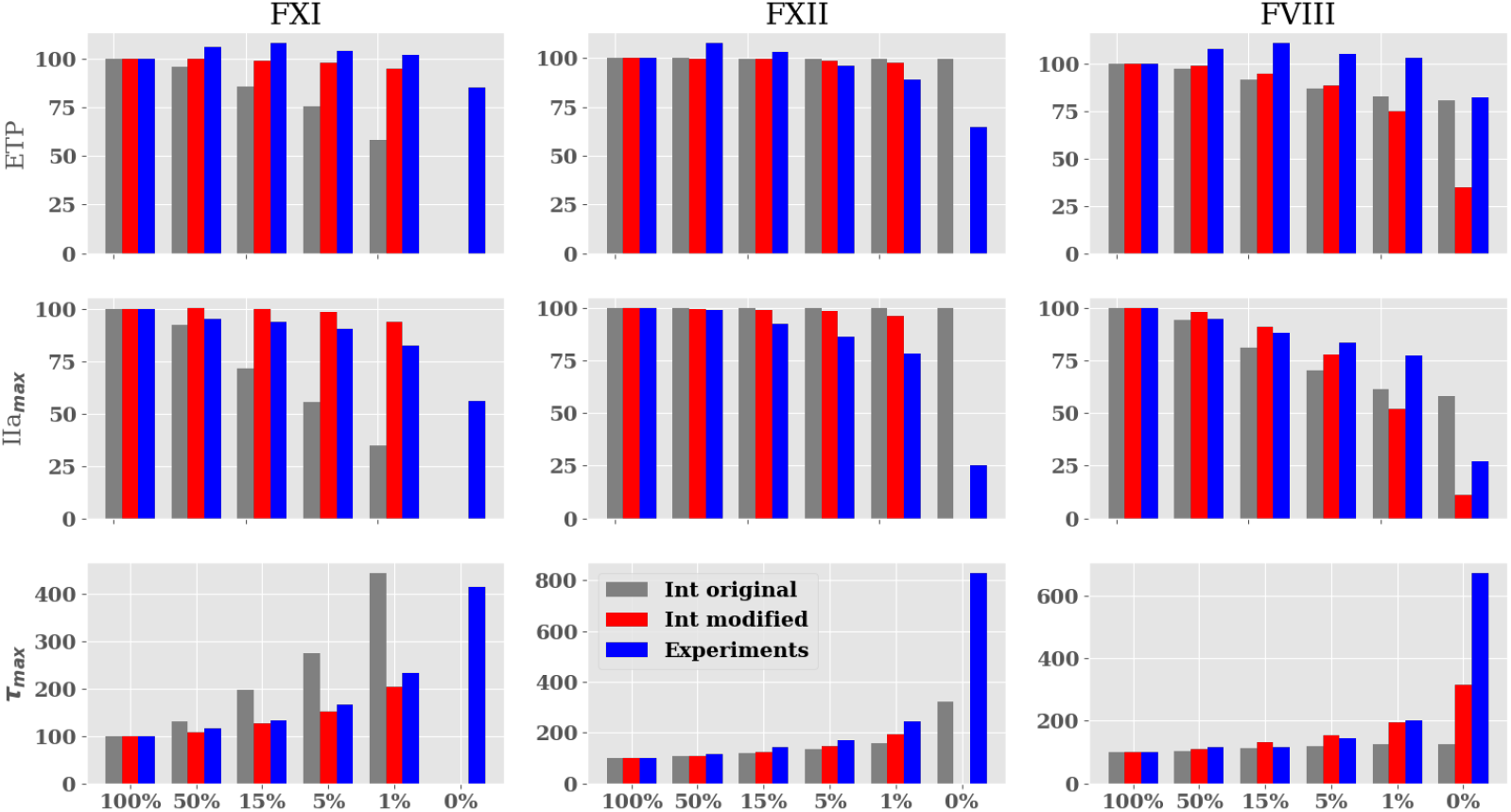
Evolution of the different parameters (%) for each factor concentration range (%) after contact activation. 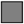 *In silico* TG obtained by Int original; 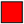 obtained by Int modified; 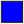 experimental results. In the 0% experimental case for FXII and FXI thrombin is produced since residual amounts of coagulation factors are still present in plasma.

First, *in silico* thrombin formation for FXII the 0% concentration should not take place since factor FXII activation is the starting point of the coagulation case. The reason for normal, yet delayed, thrombin production at 0% concentration relies on the activation of FX and FIX by activated FVII (FVIIa), independently of TF, by reactions:

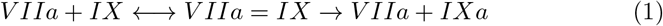

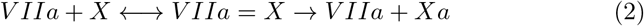

This mechanism was originally proposed by Komiyama *et al.* [23] and incorporated in the Int model. Komiyama *et al.* used 1% of the total FVII concentration in its activated state in order to trigger the reactions and characterize the kinetic rates. The trouble with this rationale is that there are at least three forms of FVII: (i) FVII zymogen, (ii) zymogen-like FVII which corresponds to the 1% circulating activated free form of FVII and (iii) activated FVII which is either bound to TF under physiological conditions or unbound in therapeutic contexts corresponding to the recombinant FVIIa (Novoseven^®^, Novo Nordisk A/S, Danemark). Reactions (1) and (2) do not reflect physiological mechanisms in the absence of circulating TF, but describe the non-physiological recombinant FVIIa (Novoseven^®^) interactions [23]. Therefore, both reactions (1) and (2) were deleted in both the original and modified Int models. Figure 3 shows that by omitting reactions (1) and (2) numerical thrombin formation at the 0% FXII case is suppressed.

The activation of FXII due to STA^®^-PTT-A^®^ reagent was set to *k*_1_ = 5 × 10^3^. Since the original value of Chatterjee *et al.* [2] was fitted to their experimental data, therefore, it is not universal for FXII contact activation. This arbitrary increase of the kinetic rate was motivated to reduce the *τ_max_* value which as shown in Fig. 2 lags the experimental data.

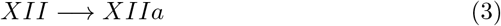

The third major modification to the Int model is related to FXI activation mechanisms. In the original model, FXI is activated by FXIIa, thrombin and an auto-activation process, respectively:

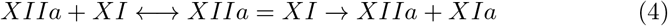

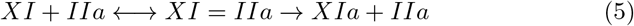

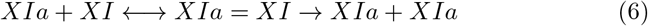

*In vitro* FXI activation by thrombin is mainly reported with circulated platelets [24] and inhibited by fibrinogen [25]. As TGA experiments were performed on platelet poor plasma (PPP), reaction (5) was deleted from the model. The work of Gailani and Bronze [25] is referenced in the original Int model for reaction (4). Gailani and Bronze [25] stated that the kinetic constants were not determined with precision because it exceeded the achievable factor concentration. Thus, the kinetic constants involved in FXI activation by FXIIa were modified to improve the fit with experimental data (*K*_1_ = 7 × 10^8^ M^−1^ s^−1^, *K*_−1_ = 200 s^−1^ *k_cat_* = 0.002 s^−1^). With the same rationale, the kinetic constant of reaction (6) describing the FXI auto-activation was reduced to an optimal value of *K*_1_ = 0.7975 × 10^6^ M^−1^ s^−1^ 4-fold lower than the value reported by Kramoroff *et al.* [24].

Additional modifications were done regarding the cofactor function of FVI-IIa. *In silico* thrombin generation parameters showed a larger pro-coagulant profile compared to *in vitro* data, especially for concentrations of FVIII under 1% (see Figures 2 and 4), suggesting that the numerical model overestimated the enzymatic activity of FIXa controlled by reactions (7) and (8).

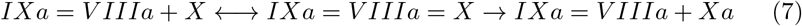

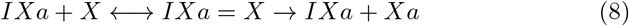

**Fig. 4.**
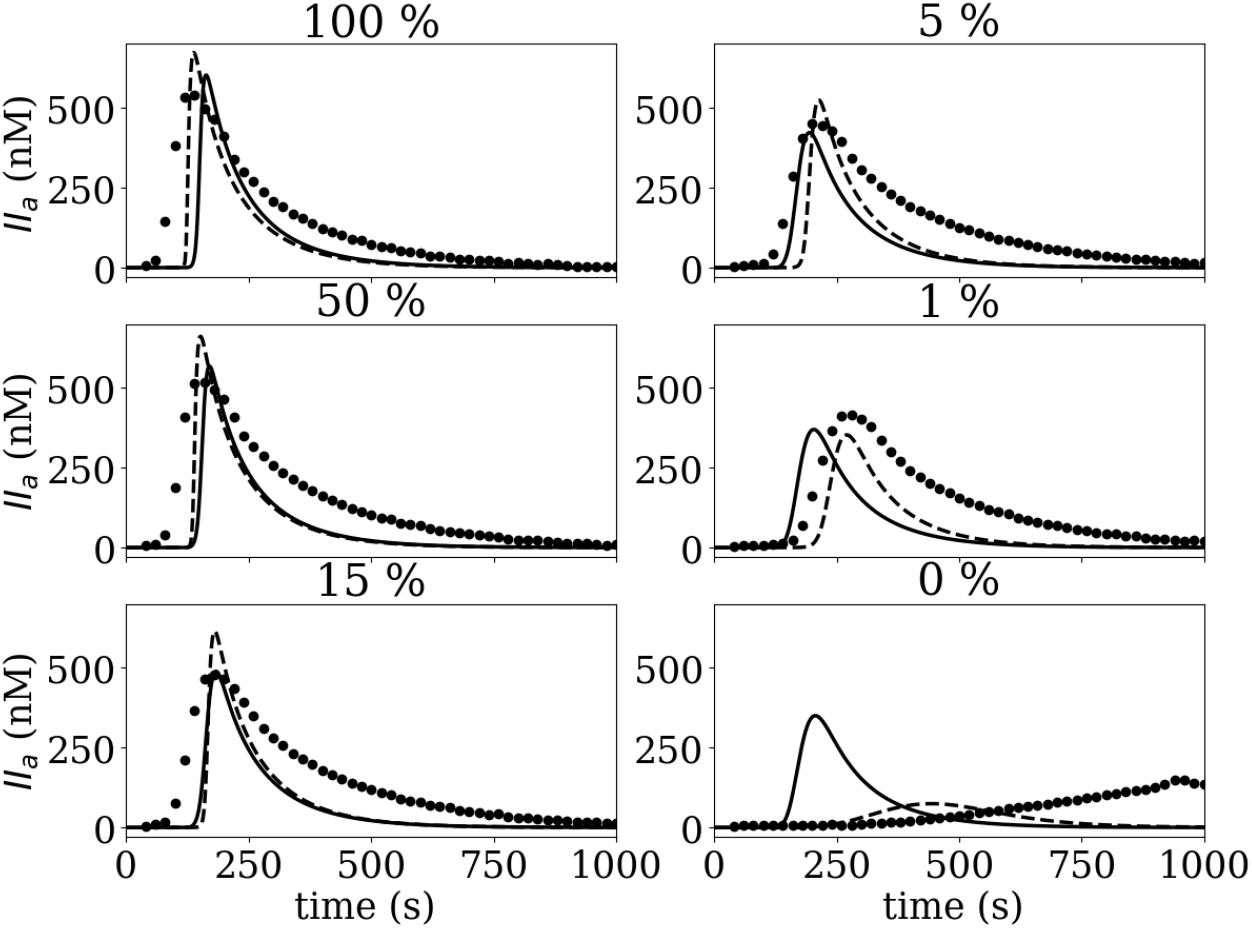
Thrombin generation (TG) for FVIII concentration range after contact activation the solid line - is the *in silico* data obtained by Int original; the dashed line -- is obtained by Int modified, and • is the experimental data.

The kinetic constants of reaction (7) and (8) were modified to decrease the activation of the FX by the unbound FIXa and the intrinsic tenase complex FIXa-FVIIIa. The kinetic constants were obtained from Kogan *et al.* [26] who characterized this reaction focusing on the contact system. The aforementioned modifications improved the numerical model predictions of ETP, IIa_*max*_, and *τ_max_* for the three ranges of coagulation factors concentrations as observed in Fig. 3. The only cases in which modifications reduced the accuracy of the numerical model were the ETP and IIa_*max*_ values for FVIII. To understand better the underlying behavior in the FVIII case, thrombin production is presented in Fig. 4. Figure 4 shows the thrombin production curves for the TGA data, Int original, and Int modified numerical data. A reduction of the pro-coagulant profile can be observed in the modified data, especially for the 1 and 0% cases. In addition, *τ_max_* has been increased following the trend observed experimentally. The thrombin formation trend for the modified model appears to have a better agreement with experimental data suggesting that the discrepancy observed in ETP and IIa_*max*_ values of Fig. 3 are not indicative of the actual thrombin formation dynamics at low concentrations.

### 3.2 Ext model for the TF pathway

Figure 5 compares the original Ext model thrombin formation to experimental TGA considering physiological plasma. The numerical results show a delayed and reduced thrombin formation compared to the experimental data. This is in line with other numerical studies [14] which reported a thrombin peak formation after 1600 seconds for a physiological plasma sample. In contrast, our experimental TGA thrombin formation peaks around 550 seconds.

**Fig. 5.**
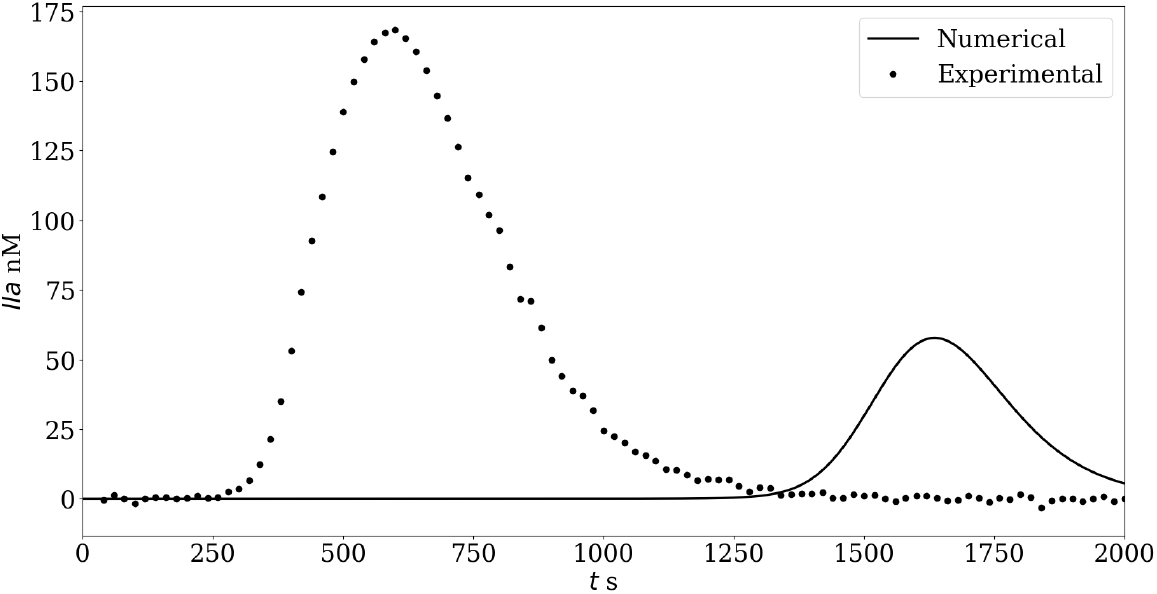
Thrombin formation in a physiological plasma sample. The original Ext model shows a low and delayed thrombin generation.

A possible explanation for this observation is that the FVIIa-TF complex formation is independent of calcium concentration and may occur *in vitro* during the TGA pre-incubation phase [27]. In that case, at the beginning of the experimental TGA, the calcium can immediately lead to the activation of FX through the FVIIa-TF complex. To verify this hypothesis, the FVIIa-TF complex concentration was changed *in silico*, modifying the initial concentrations of FVIIa-TF and TF from 0 pM/1 pM to 1 pM/0 pM in both in-silico models. The former values correspond to the nominal value (see Table1) while the latter correspond to the case where all the TF available has been consumed to form the FVIIa-TF complex before the injection of Calcium at the start of the experiment.

Thrombin formation curves for the original and modified Ext models, as well as experimental TGA data, are displayed in Fig. 6 for a range of FVIII concentrations. Remarkable agreement between the numerical and experimental data is observed when the initial value of the FVIIa-TF complex was set to 1 pM.

**Fig. 6.**
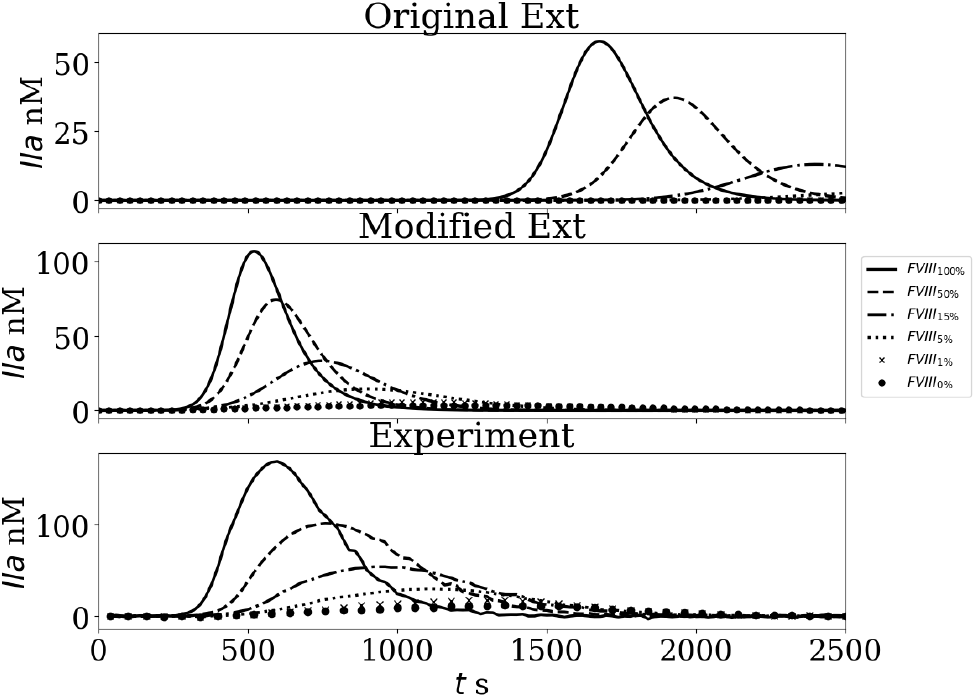
Thrombin generation (TG) for FVIII concentration range after TF activation A: *In silico* TG obtained by the original Ext; B: obtained by modified Ext; C: experimental curves.

Figure 7 shows the relative values of ETP, *IIa_max_*, and *τ_max_* for the original model setting the concentration of FVIIa-TF complex to 1pM, the modified Ext model and experimental data. Note that the agreement between the original model with FVIIa-TF = 1 pM and the experimental values is not good for very small FVIII concentrations (≤ 1%). This observation could be explained by an overestimation of FVIII activity in the Ext model. To explore this hypothesis and improve the activity of the uncomplexed FIXa, the kinetic constants of Eq. (8) were modified. An arbitrarily 60 fold increase of the catalytic constant (Kcat = 4.8 × 10^−2^ s^−1^ instead of Kcat = 8.0 × 10^−4^ s^−1^ from [3]). The modified Ext model improved the results remarkably for ETP, IIa_*max*_ and *τ_max_* variables for the 5, 1, and 0% concentrations.

**Fig. 7.**
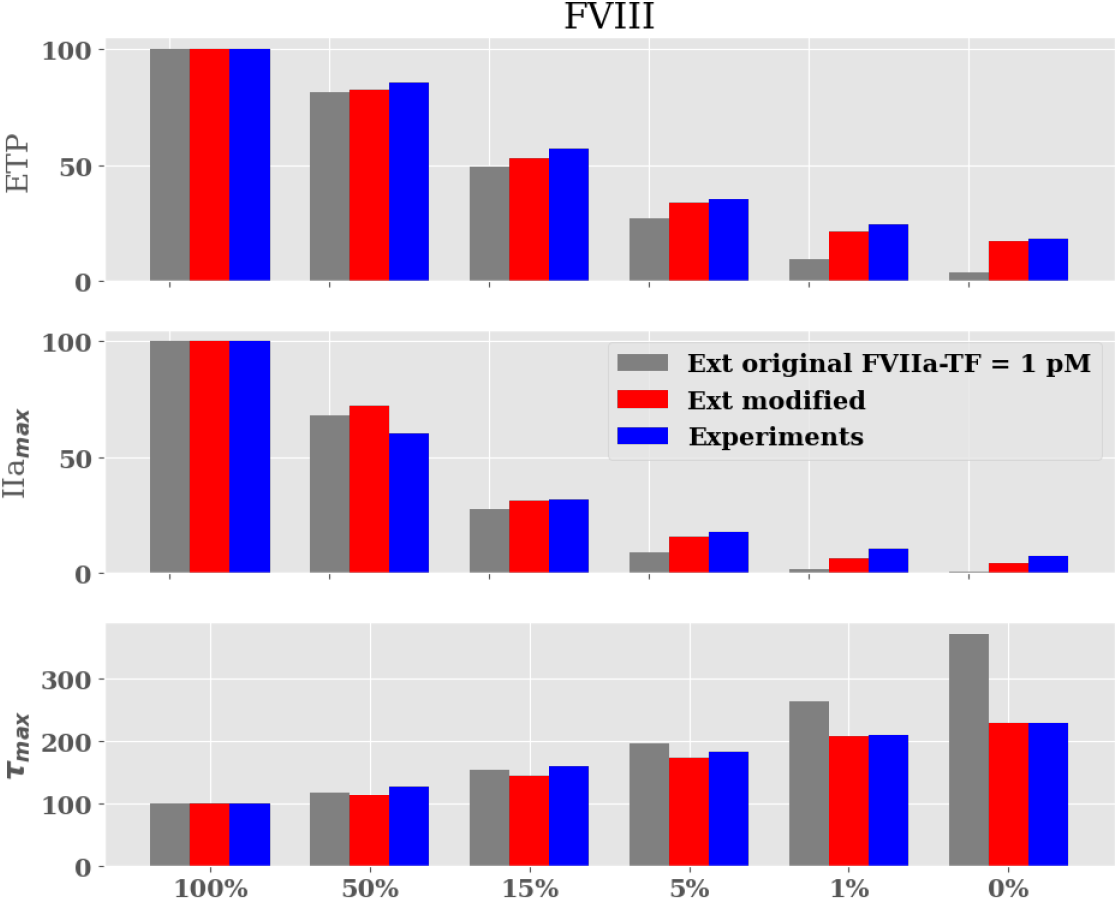
Evolution of the ETP 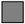 *In silico* ETP obtained by the original Ext model with the initial concentration FVIIa-TF complex = 1pM; 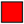 obtained by modified Ext model; 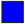 experimental results.

## 4 Discussion and Conclusion

In recent studies, mathematical models of the coagulation cascade have been used to shed light on mechanisms leading to clotting disorders. Link *et al.* [29, 8] performed a global sensitivity analysis using the coagulation model of Kuharsky and Fogelson [30] that led to the identification of FV as a key modifier of thrombin generation among hemophilia A patients. Their results were confirmed later with additional experimental assays highlighting the value of using the numerical model to build a synthetic data set to construct and test specific hypotheses. Brummel-Ziedins *et al.* [31] developed a FVII deficiency model and Anand *et al.* [32] studied protein C and antithrombin deficiencies by comparing their numerical results to thrombin generation experiments initiated by TF-VIIa [33]. Models have also been used to study new treatment strategies that would be further assessed *in vitro*. For instance, Burghaus *et al.* [5] and Brummel-Ziedins *et al.* [34] investigated the rivaroxaban effects. Adams *et al.* [35] and recently Zavialova *et al.* [36] worked on the possible outcomes of direct thrombin inhibitors. Despite great progress in thrombin formation simulations, a direct comparison between numerical models and *in vitro* experiments is frequently inappropriate since the mathematical models are constructed for specific and ideal conditions as pointed out by Link *et al.* [12,8]. Chelle *et al.* [14] tested the performance of five coagulation models by comparing their results under relevant clinical scenarios. They found that the predictive capabilities of the models were far from acceptable and needed an additional optimization step using genetic algorithms to adapt kinetic parameters and reproduce *in vitro* data. In particular, they suggested that performing patient-specific optimization is needed in order to achieve good model estimations. However, if each new application requires patient-specific calibration, the global functionality of the model is reduced. A strategy for improving the model instead of performing complex optimization work is to evaluate the sensitivity of the model to a parameter that represents a clinical or pathological situation. In this way, not only the actual value of a given reaction rate is tested but also the intrinsic interactions of coagulation reactions and their response to a change in a coagulation factor concentration.

In this study, two well known models of the coagulation cascade were compared with TGA experimental results and proper modifications are suggested to improve their accuracy. The modifications of the Int model for the contact phase are related to excluding the reactions related to spurious delayed thrombin formation due to FVIIa activation in the absence of TF. In addition, FXI activation by thrombin was not considered since platelet poor plasma was used in our study. Then, kinetic constants involved in FXI auto-activation and activation by XIIa were fitted to improve the comparison with TGA results.

Another modification regarding excessive procoagulant activity by FVIIIa is proposed to reduce the activation of FX by unbounded FIXa and IXa=VIIIa. As pointed out in [24, 25] the values of the kinetic rates are not very precise and thus are susceptible to change. Concerning the Ext model, the main consideration was to introduce 1 pM initial concentration of FVIIa-TF, which is motivated by the formation of complex FVIIa-TF during the incubation phase of the TGA experiment. This adjustment allowed to shorten the delay of thrombin formation. The modification on the Int and Ext models improved the comparison for ETP, IIa_*max*_ and *τ_max_* overall. It is noteworthy that even though the comparisons of the original model to the 100% ‘nominal’ case are fairly good, once the models are evaluated under pathological conditions of FXII, FXI and FVIII deficiencies, the modifications improved the comparison for the three variables observed and prevented artificial thrombin formation for the 0% concentration cases. Note however that, in the case of FVIII (< 1%) triggered via the Intrinsic pathway, both the original and modified Int models fail to reproduce the ETP, IIa_*max*_ and *τ_max_.* Therefore, improvements to the model should be further considered when looking at severe hemophilic patients.

The proposed modifications to the Int and Ext models are not universal (as shown in reaction 8 with different values for Int and Ext models) but are well suited for the TGA experimental conditions studied. Thus, they can be used as a valuable tool to explore any scenario that might be costly experimentally. Since the coagulation model still has limitations linked to modeling assumptions, careful consideration should be taken when extrapolating the models to open system applications such as thrombosis simulations of medical devices or organ scale studies. Considering the complexity of kinetic schemes of the coagulation cascade, the authors recommend close collaboration between hematologists and modelers when applying coagulation models. Continuous validation of coagulation models in complex situations such as pathological cases is needed in order to broaden their application scope. This remains a necessary effort towards robust and reliable coagulation models that could be used to study coagulation disorders and in drug or bio-medical devices development.

## Acknowledgements

The authors would like to thank Prof. Jean-François Schved for his insightful comments. This research was supported by CONACyT, Mexico scholarship and the LabEx Numev (convention ANR-10-LABX-20). This work was publicly funded through ANR (the French National Research Agency) under the “Investissements d’avenir” programme with the reference ANR-16-IDEX-0006.

## Conflict of interest

The authors declare that they have no conflict of interest.

## Footnotes

https://imag.umontpellier.fr/~yales2bio/index.html

## Notes

### Competing Interest Statement

The authors have declared no competing interest.

