## Supplemental Table 1, 2, 3, and 4 for "Critical evaluation of kinetic schemes for coagulation"

SUPPLEMENTARY INFORMATION

**Table 1. Original Int Model from Chatterjee et al. [2]**

| # | Reaction | $k_1$ | $k_{-1}$ | $k_{cat}$ |
|---|---|---|---|---|
| 1 | TF + VII ↔ TF = VII | $3.2x10^6$ $M^{-1}s^{-1}$ | $3.1x10^{-3}$ $s^{-1}$ | |
| 2 | TF + VIIa ↔ TF = VIIa | $2.3x10^7$ $M^{-1}s^{-1}$ | $3.1x10^{-3}$ $s^{-1}$ | |
| 3 | TF = VIIa + VII → TF = VIIa + VIIa | $4.4x10^5$ $M^{-1}s^{-1}$ | | |
| 4 | Xa + VII → Xa + VIIa | $1.3x10^7$ $M^{-1}s^{-1}$ | | |
| 5 | IIa + VII → IIa + VIIa | $2.3x10^4$ $M^{-1}s^{-1}$ | | |
| 6 | TF = VIIa + X ↔ TF = VIIa = X → TF = VIIa = Xa | $2.5x10^7$ $M^{-1}s^{-1}$ | 1.05 $s^{-1}$ | 6 $s^{-1}$ |
| 7 | TF = VIIa + Xa ↔ TF = VIIa = Xa | $2.2x10^7$ $M^{-1}s^{-1}$ | 19 $s^{-1}$ | |
| 8 | TF = VIIa + IX ↔ TF = VIIa = IX → TF = VIIa + IXa | $1.0x10^7$ $M^{-1}s^{-1}$ | 2.4 $s^{-1}$ | 1.8 $s^{-1}$ |
| 9 | II + Xa → IIa + Xa | $7.5x10^3$ $M^{-1}s^{-1}$ | | |
| 10 | IIa + VIII → IIa + VIIIa | $2.0x10^7$ $M^{-1}s^{-1}$ | | |
| 11 | VIIIa + IXa ↔ IXa = VIIIa | $1.0x10^7$ $M^{-1}s^{-1}$ | $5.0x10^{-3}$ $s^{-1}$ | |
| 12 | IXa = VIIIa + X ↔ IXa = VIIIa = X → IXa = VIIIa + Xa | $1.0x10^8$ $M^{-1}s^{-1}$ | $1.0x10^{-3}$ $s^{-1}$ | 8.2 $s^{-1}$ |
| 13 | VIIIa ↔ $VIIIa_1$ + $VIIIa_2$ | $6.0x10^{-3}$ $s^{-1}$ | $2.2x10^4$ $M^{-1}s^{-1}$ | |
| 14 | IXa = VIIIa = X → IXa + X + $VIIIa_1$ + $VIIIa_2$ | $1.0x10^{-3}$ $s^{-1}$ | | |
| 15 | IXa = VIIIa → IXa + $VIIIa_1$ + $VIIIa_2$ | $1.0x10^{-3}$ $s^{-1}$ | | |
| 16 | IIa + V → IIa + Va | $2.0x10^7$ $M^{-1}s^{-1}$ | | |
| 17 | Xa + Va ↔ Xa = Va | $4.0x10^8$ $M^{-1}s^{-1}$ | 0.2 $s^{-1}$ | |
| 18 | Xa = Va + II ↔ Xa = Va = II → Xa = Va + mIIa | $1.0x10^8$ $M^{-1}s^{-1}$ | 103 $s^{-1}$ | 63.5 $s^{-1}$ |
| 19 | Xa = Va + mIIa → Xa = Va + IIa | $1.5x10^7$ $M^{-1}s^{-1}$ | | |
| 20 | Xa + TFPI ↔ Xa = TFPI | $9.0x10^5$ $M^{-1}s^{-1}$ | $3.6x10^{-4}$ $s^{-1}$ | |
| 21 | TF = VIIa = Xa + TFPI ↔ TF = VIIa = Xa = TFPI | $3.2x10^8$ $M^{-1}s^{-1}$ | $1.1x10^{-4}$ $s^{-1}$ | |
| 22 | TF = VIIa + Xa = TFPI → TF = VIIa = Xa = TFPI | $5.0x10^7$ $M^{-1}s^{-1}$ | | |
| 23 | Xa + ATIII → Xa = ATIII | $1.5x10^3$ $M^{-1}s^{-1}$ | | |
| 24 | mIIa + ATIII → mIIa = ATIII | $7.1x10^3$ $M^{-1}s^{-1}$ | | |
| 25 | IXa + ATIII → IXa = ATIII | $4.9x10^2$ $M^{-1}s^{-1}$ | | |
| 26 | IIa + ATIII → IIa = ATIII | $7.1x10^3$ $M^{-1}s^{-1}$ | | |
| 27 | TF = VIIa + ATIII → TF = VIIa = ATIII | $2.3x10^2$ $M^{-1}s^{-1}$ | | |

| | | | | |
|---|---|---|---|---|
| 28 | Boc-VPR-MCA + IIa ↔ Boc-VPR-MCA = IIa → Boc-VPR + MCA + IIa | 1.0x10$^{8}$ M$^{-1}$s$^{-1}$ | 6.1x10$^{3}$ s$^{-1}$ | 53.8 s$^{-1}$ |
| 29 | XII → XIIa | 5.0x10$^{-4}$ s$^{-1}$ | | 3.3x10$^{-2}$ s$^{-1}$ |
| 30 | XIIa + XII ↔ XIIa = XII → XIIa + XIIa | 1.0x10$^{8}$ M$^{-1}$s$^{-1}$ | 750 s$^{-1}$ | 40 s$^{-1}$ |
| 31 | XIIa + PK ↔ XIIa = PK → XIIa + K | 1.0x10$^{8}$ M$^{-1}$s$^{-1}$ | 3.6x10$^{3}$ s$^{-1}$ | 5.7 s$^{-1}$ |
| 32 | XII + K ↔ XII = K → XIIa + K | 1.0x10$^{8}$ M$^{-1}$s$^{-1}$ | 45.3 s$^{-1}$ | |
| 33 | PK + K → K + K | 2.7x10$^{4}$ M$^{-1}$s$^{-1}$ | | |
| 34 | K → K.inhibited | 1.1x10$^{-2}$ s$^{-1}$ | | |
| ~~35~~ | ~~XIIa + CTI ↔ XIIa = CTI~~ | ~~1.0x10$^{8}$ M$^{-1}$s$^{-1}$~~ | ~~2.4 s$^{-1}$~~ | |
| 36 | XIIa + C1inh → XIIa = C1inh | 3.6x10$^{3}$ M$^{-1}$s$^{-1}$ | | |
| 37 | XIIa + ATIII → XIIa = ATIII | 21.6 M$^{-1}$s$^{-1}$ | | |
| 38 | XI + IIa ↔ XI = IIa → XIa + IIa | 1.0x10$^{8}$ M$^{-1}$s$^{-1}$ | 5 s$^{-1}$ | 1.3x10$^{-4}$ s$^{-1}$ |
| 39 | XIIa + XI ↔ XIIa = XI → XIIa + XIa | 1.0x10$^{8}$ M$^{-1}$s$^{-1}$ | 200 s$^{-1}$ | 5.7x10$^{-4}$ s$^{-1}$ |
| 40 | XIa + XI ↔ XIa = XI→ XIa + XIa | 3.19x10$^{6}$ M$^{-1}$s$^{-1}$ | | |
| 41 | XIa + ATIII → XIa = ATIII | 3.2x10$^{2}$ M$^{-1}$s$^{-1}$ | | |
| 42 | XIa + C1inh → XIa = C1inh | 1.8x10$^{3}$ M$^{-1}$s$^{-1}$ | | |
| 43 | XIa + α1AT → XIa = α1AT | 1.0x10$^{2}$ M$^{-1}$s$^{-1}$ | | |
| 44 | XIa + α2AP → XIa = α2AP | 4.3x10$^{3}$ M$^{-1}$s$^{-1}$ | | |
| 45 | XIa + IX ↔ XIa = IX → XIa + IXa | 1.0x10$^{8}$ M$^{-1}$s$^{-1}$ | 41 s$^{-1}$ | 7.7 s$^{-1}$ |
| 46 | IXa + X ↔ IXa = X → IXa + Xa | 1.0x10$^{8}$ M$^{-1}$s$^{-1}$ | 0.64 s$^{-1}$ | 7.0x10$^{-4}$ s$^{-1}$ |
| 47 | Xa + VIII ↔ Xa = VIII → Xa + VIIIa | 1.0x10$^{8}$ M$^{-1}$s$^{-1}$ | 2.1 s$^{-1}$ | 0.023 s$^{-1}$ |
| 48 | VIIa + IX ↔ VIIa = IX → VIIa + IXa | 1.0x10$^{8}$ M$^{-1}$s$^{-1}$ | 0.9 s$^{-1}$ | 3.6x10$^{-5}$ s$^{-1}$ |
| 49 | VIIa + X ↔ VIIa = X → VIIa + Xa | 1.0x10$^{8}$ M$^{-1}$s$^{-1}$ | 210 s$^{-1}$ | 1.6x10$^{-6}$ s$^{-1}$ |
| ~~50~~ | ~~Fbg + IIa ↔ Fbg = IIa → Fbn1 + IIa + FPA~~ | ~~1.0x10$^{8}$ M$^{-1}$s$^{-1}$~~ | ~~636 s$^{-1}$~~ | ~~84 s$^{-1}$~~ |
| ~~51~~ | ~~Fbn1 + IIa ↔ Fbn1 = IIa → Fbn2 + IIa + FPB~~ | ~~1.0x10$^{8}$ M$^{-1}$s$^{-1}$~~ | ~~742.6 s$^{-1}$~~ | ~~7.4 s$^{-1}$~~ |
| ~~52~~ | ~~2Fbn1 ↔ (Fbn1)$_2$~~ | ~~1.0x10$^{6}$ M$^{-1}$s$^{-1}$~~ | ~~6.4 x10$^{-2}$ s$^{-1}$~~ | |
| ~~53~~ | ~~(Fbn1)$_2$ + IIa ↔ (Fbn1)$_2$ = IIa → (Fbn2)$_2$ + IIa + FPB~~ | ~~1.0x10$^{8}$ M$^{-1}$s$^{-1}$~~ | ~~701 s$^{-1}$~~ | ~~49 s$^{-1}$~~ |
| ~~54~~ | ~~Fbn2 + IIa ↔ Fbn2 = IIa~~ | ~~1.0x10$^{8}$ M$^{-1}$s$^{-1}$~~ | ~~1.0 x10$^{3}$ s$^{-1}$~~ | |
| ~~55~~ | ~~(Fbn1)$_2$ = IIa + ATIII → (Fbn1)$_2$ = IIa = ATIII~~ | ~~1.6x10$^{4}$ M$^{-1}$s$^{-1}$~~ | | |
| ~~56~~ | ~~Fbn1 = IIa + ATIII → Fbn1 = IIa = ATIII~~ | ~~1.6x10$^{4}$ M$^{-1}$s$^{-1}$~~ | | |
| ~~57~~ | ~~Fbn2 = IIa + ATIII → Fbn2 = IIa = ATIII~~ | ~~1.6x10$^{4}$ M$^{-1}$s$^{-1}$~~ | | |

**Table 2. Original Ext Model from Hocking et al. [1] and Butenas et al. [3]**

| # | Reaction | $k_1$ | $k_{-1}$ | $k_{cat}$ |
|---|---|---|---|---|
| 1 | TF + VII ↔ TF = VII | $3.2 \times 10^6$ $M^{-1}s^{-1}$ | $3.1 \times 10^{-3}$ $s^{-1}$ | |
| 2 | TF + VIIa ↔ TF = VIIa | $2.3 \times 10^7$ $M^{-1}s^{-1}$ | $3.1 \times 10^{-3}$ $s^{-1}$ | |
| 3 | TF = VIIa + VII → TF = VIIa + VIIa | $4.4 \times 10^5$ $M^{-1}s^{-1}$ | | |
| 4 | Xa + VII → Xa + VIIa | $1.3 \times 10^7$ $M^{-1}s^{-1}$ | | |
| 5 | IIa + VII → IIa + VIIa | $2.3 \times 10^4$ $M^{-1}s^{-1}$ | | |
| 6 | TF = VIIa + X ↔ TF = VIIa = X → TF = VIIa = Xa | $2.5 \times 10^7$ $M^{-1}s^{-1}$ | 1.05 $s^{-1}$ | 6 $s^{-1}$ |
| 7 | TF = VIIa + Xa ↔ TF = VIIa = Xa | $2.2 \times 10^7$ $M^{-1}s^{-1}$ | 19 $s^{-1}$ | |
| 8 | TF = VIIa + IX ↔ TF = VIIa = IX → TF = VIIa + IXa | $1.0 \times 10^7$ $M^{-1}s^{-1}$ | 2.4 $s^{-1}$ | 1.8 $s^{-1}$ |
| 9 | II + Xa → IIa + Xa | $7.5 \times 10^3$ $M^{-1}s^{-1}$ | | |
| 10 | IIa + VIII → IIa + VIIIa | $2.0 \times 10^7$ $M^{-1}s^{-1}$ | | |
| 11 | VIIIa + IXa ↔ IXa = VIIIa | $1.0 \times 10^7$ $M^{-1}s^{-1}$ | $5.0 \times 10^{-3}$ $s^{-1}$ | |
| 12 | IXa = VIIIa + X ↔ IXa = VIIIa = X → IXa = VIIIa + Xa | $1.0 \times 10^8$ $M^{-1}s^{-1}$ | $1.0 \times 10^{-3}$ $s^{-1}$ | 8.2 $s^{-1}$ |
| 13 | VIIIa ↔ $VIIIa_1$ + $VIIIa_2$ | $6.0 \times 10^{-3}$ $s^{-1}$ | $2.2 \times 10^4$ $M^{-1}s^{-1}$ | |
| 14 | IXa = VIIIa = X → IXa + X + $VIIIa_1$ + $VIIIa_2$ | $1.0 \times 10^{-3}$ $s^{-1}$ | | |
| 15 | IXa = VIIIa → IXa + $VIIIa_1$ + $VIIIa_2$ | $1.0 \times 10^{-3}$ $s^{-1}$ | | |
| 16 | IIa + V → IIa + Va | $2.0 \times 10^7$ $M^{-1}s^{-1}$ | | |
| 17 | Xa + Va ↔ Xa = Va | $4.0 \times 10^8$ $M^{-1}s^{-1}$ | 0.2 $s^{-1}$ | |
| 18 | Xa = Va + II ↔ Xa = Va = II → Xa = Va + mIIa | $1.0 \times 10^8$ $M^{-1}s^{-1}$ | 103 $s^{-1}$ | 63.5 $s^{-1}$ |
| 19 | Xa = Va + mIIa → Xa = Va + IIa | $1.5 \times 10^7$ $M^{-1}s^{-1}$ | | |
| 20 | Xa + TFPI ↔ Xa = TFPI | $9.0 \times 10^5$ $M^{-1}s^{-1}$ | $3.6 \times 10^{-4}$ $s^{-1}$ | |
| 21 | TF = VIIa = Xa + TFPI ↔ TF = VIIa = Xa = TFPI | $3.2 \times 10^8$ $M^{-1}s^{-1}$ | $1.1 \times 10^{-4}$ $s^{-1}$ | |
| 22 | TF = VIIa + Xa = TFPI → TF = VIIa = Xa + TFPI | $5.0 \times 10^7$ $M^{-1}s^{-1}$ | | |
| 23 | Xa + ATIII → Xa = ATIII | $1.5 \times 10^3$ $M^{-1}s^{-1}$ | | |
| 24 | mIIa + ATIII → mIIa = ATIII | $7.1 \times 10^3$ $M^{-1}s^{-1}$ | | |
| 25 | IXa + ATIII → IXa = ATIII | $4.9 \times 10^2$ $M^{-1}s^{-1}$ | | |
| 26 | IIa + ATIII → IIa = ATIII | $7.1 \times 10^3$ $M^{-1}s^{-1}$ | | |
| 27 | TF = VIIa + ATIII → TF = VIIa = ATIII | $2.3 \times 10^2$ $M^{-1}s^{-1}$ | | |

| | | | |
|---|---|---|---|
| 28 | IXa + X → IXa + Xa | $K_m$ = 1,4x10$^{-7}$ M | $K_{cat}$ = 8x10$^{-4}$ $s^{-1}$ |

| **Table 3. Modified Int Model** | | | | |
|---|---|---|---|---|
| # | Reaction | $k_1$ | $k_{-1}$ | $k_{cat}$ |
| 1 | TF + VII ↔ TF = VII | $3.2x10^6\ M^{-1}s^{-1}$ | $3.1x10^{-3}\ s^{-1}$ | |
| 2 | TF + VIIa ↔ TF = VIIa | $2.3x10^7\ M^{-1}s^{-1}$ | $3.1x10^{-3}\ s^{-1}$ | |
| 3 | TF = VIIa + VII ➔ TF = VIIa + VIIa | $4.4x10^5\ M^{-1}s^{-1}$ | | |
| 4 | Xa + VII ➔ Xa + VIIa | $1.3x10^7\ M^{-1}s^{-1}$ | | |
| 5 | IIa + VII ➔ IIa + VIIa | $2.3x10^4\ M^{-1}s^{-1}$ | | |
| 6 | TF = VIIa + X ↔ TF = VIIa = X ➔ TF = VIIa = Xa | $2.5x10^7\ M^{-1}s^{-1}$ | $1.05\ s^{-1}$ | $6\ s^{-1}$ |
| 7 | TF = VIIa + Xa ↔ TF = VIIa = Xa | $2.2x10^7\ M^{-1}s^{-1}$ | $19\ s^{-1}$ | |
| 8 | TF = VIIa + IX ↔ TF = VIIa = IX ➔ TF = VIIa + IXa | $1.0x10^7\ M^{-1}s^{-1}$ | $2.4\ s^{-1}$ | $1.8\ s^{-1}$ |
| 9 | II + Xa ➔ IIa + Xa | $7.5x10^3\ M^{-1}s^{-1}$ | | |
| 10 | IIa + VIII ➔ IIa + VIIIa | $2.0x10^7\ M^{-1}s^{-1}$ | | |
| 11 | VIIIa + IXa ↔ IXa = VIIIa | $1.0x10^7\ M^{-1}s^{-1}$ | $5.0x10^{-3}\ s^{-1}$ | |
| 12 | IXa = VIIIa + X ➔ IXa = VIIIa + Xa | $0.19x10^{-6}\ M$ | | $29\ s^{-1}$ |
| 13 | VIIIa ↔ $VIIIa_1$ + $VIIIa_2$ | $6.0x10^{-3}\ s^{-1}$ | $2.2x10^4\ M^{-1}s^{-1}$ | |
| 14 | IXa = VIIIa = X ➔ IXa + X + $VIIIa_1$ + $VIIIa_2$ | $1.0x10^{-3}\ s^{-1}$ | | |
| 15 | IXa = VIIIa ➔ IXa + $VIIIa_1$ + $VIIIa_2$ | $1.0x10^{-3}\ s^{-1}$ | | |
| 16 | IIa + V ➔ IIa + Va | $2.0x10^7\ M^{-1}s^{-1}$ | | |
| 17 | Xa + Va ↔ Xa = Va | $4.0x10^8\ M^{-1}s^{-1}$ | $0.2\ s^{-1}$ | |
| 18 | Xa = Va + II ↔ Xa = Va = II ➔ Xa = Va + mIIa | $1.0x10^8\ M^{-1}s^{-1}$ | $103\ s^{-1}$ | $63.5\ s^{-1}$ |
| 19 | Xa = Va + mIIa ➔ Xa = Va + IIa | $1.5x10^7\ M^{-1}s^{-1}$ | | |
| 20 | Xa + TFPI ↔ Xa = TFPI | $9.0x10^5\ M^{-1}s^{-1}$ | $3.6x10^{-4}\ s^{-1}$ | |
| 21 | TF = VIIa = Xa + TFPI ↔ TF = VIIa = Xa = TFPI | $3.2x10^8\ M^{-1}s^{-1}$ | $1.1x10^{-4}\ s^{-1}$ | |
| 22 | TF = VIIa + Xa = TFPI ➔ TF = VIIa = Xa = TFPI | $5.0x10^7\ M^{-1}s^{-1}$ | | |
| 23 | Xa + ATIII ➔ Xa = ATIII | $1.5x10^3\ M^{-1}s^{-1}$ | | |
| 24 | mIIa + ATIII ➔ mIIa = ATIII | $7.1x10^3\ M^{-1}s^{-1}$ | | |
| 25 | IXa + ATIII ➔ IXa = ATIII | $4.9x10^2\ M^{-1}s^{-1}$ | | |
| 26 | IIa + ATIII ➔ IIa = ATIII | $7.1x10^3\ M^{-1}s^{-1}$ | | |
| 27 | TF = VIIa + ATIII ➔ TF = VIIa = ATIII | $2.3x10^2\ M^{-1}s^{-1}$ | | |
| 28 | Boc-VPR-MCA + IIa ↔ Boc-VPR-MCA = IIa ➔ Boc- | $1.0x10^8\ M^{-1}s^{-1}$ | $6.1x10^3\ s^{-1}$ | $53.8\ s^{-1}$ |

| | | | | |
|---|---|---|---|---|
| | VPR + MCA + IIa | | | |
| 29 | XII ➔ XIIa | $5.0x10^{-3}$ $s^{-1}$ | | |
| 30 | XIIa + XII ↔ XIIa = XII ➔ XIIa + XIIa | $1.0x10^{8}$ $M^{-1}s^{-1}$ | 750 $s^{-1}$ | $3.3x10^{-2}$ $s^{-1}$ |
| 31 | XIIa + PK ↔ XIIa = PK ➔ XIIa + K | $1.0x10^{8}$ $M^{-1}s^{-1}$ | $3.6x10^{3}$ $s^{-1}$ | 40 $s^{-1}$ |
| 32 | XII + K ↔ XII = K ➔ XIIa + K | $1.0x10^{8}$ $M^{-1}s^{-1}$ | 45.3 $s^{-1}$ | 5.7 $s^{-1}$ |
| 33 | PK + K ➔ K + K | $2.7x10^{4}$ $M^{-1}s^{-1}$ | | |
| 34 | K ➔ K.inhibited | $1.1x10^{-2}$ $s^{-1}$ | | |
| 36 | XIIa + C1inh ➔ XIIa = C1inh | $3.6x10^{3}$ $M^{-1}s^{-1}$ | | |
| 37 | XIIa + ATIII ➔ XIIa = ATIII | 21.6 $M^{-1}s^{-1}$ | | |
| 38 | XI + IIa ↔ XI = IIa ➔ XIa + IIa | $1.0x10^{8}$ $M^{-1}s^{-1}$ | 5 $s^{-1}$ | $1.3x10^{-4}$ $s^{-1}$ |
| 39 | XIIa + XI ↔ XIIa = XI ➔ XIIa + XIa | $7.0x10^{8}$ $M^{-1}s^{-1}$ | 200 $s^{-1}$ | $2x10^{-3}$ $s^{-1}$ |
| 40 | XIa + XI ↔ XIa = XI➔ XIa + XIa | $0.7975x10^{6}$ $M^{-1}s^{-1}$ | | |
| 41 | XIa + ATIII ➔ XIa = ATIII | $3.2x10^{2}$ $M^{-1}s^{-1}$ | | |
| 42 | XIa + C1inh ➔ XIa = C1inh | $1.8x10^{3}$ $M^{-1}s^{-1}$ | | |
| 43 | XIa + α1AT ➔ XIa = α1AT | $1.0x10^{2}$ $M^{-1}s^{-1}$ | | |
| 44 | XIa + α2AP ➔ XIa = α2AP | $4.3x10^{3}$ $M^{-1}s^{-1}$ | | |
| 45 | XIa + IX ↔ XIa = IX ➔ XIa + IXa | $1.0x10^{8}$ $M^{-1}s^{-1}$ | 41 $s^{-1}$ | 7.7 $s^{-1}$ |
| 46 | IXa + X ➔ IXa + Xa | $2x10^{-6}$ M | | $6.67x10^{-4}$ $s^{-1}$ |
| 47 | Xa + VIII ↔ Xa = VIII ➔ Xa + VIIIa | $1.0x10^{8}$ $M^{-1}s^{-1}$ | 2,1 $s^{-1}$ | 0.023 $s^{-1}$ |

**Table 4. Modified Ext model.**

| # | Reaction | $k_1$ | $k_{-1}$ | $k_{cat}$ |
|---|---|---|---|---|
| 1 | TF + VII ↔ TF = VII | $3.2x10^6$ $M^{-1}s^{-1}$ | $3.1x10^{-3}$ $s^{-1}$ | |
| 2 | TF + VIIa ↔ TF = VIIa | $2.3x10^7$ $M^{-1}s^{-1}$ | $3.1x10^{-3}$ $s^{-1}$ | |
| 3 | TF = VIIa + VII → TF = VIIa + VIIa | $4.4x10^5$ $M^{-1}s^{-1}$ | | |
| 4 | Xa + VII → Xa + VIIa | $1.3x10^7$ $M^{-1}s^{-1}$ | | |
| 5 | IIa + VII → IIa + VIIa | $2.3x10^4$ $M^{-1}s^{-1}$ | | |
| 6 | TF = VIIa + X ↔ TF = VIIa = X → TF = VIIa = Xa | $2.5x10^7$ $M^{-1}s^{-1}$ | 1.05 $s^{-1}$ | 6 $s^{-1}$ |
| 7 | TF = VIIa + Xa ↔ TF = VIIa = Xa | $2.2x10^7$ $M^{-1}s^{-1}$ | 19 $s^{-1}$ | |
| 8 | TF = VIIa + IX ↔ TF = VIIa = IX → TF = VIIa + IXa | $1.0x10^7$ $M^{-1}s^{-1}$ | 2.4 $s^{-1}$ | 1.8 $s^{-1}$ |
| 9 | II + Xa → IIa + Xa | $7.5x10^3$ $M^{-1}s^{-1}$ | | |
| 10 | IIa + VIII → IIa + VIIIa | $2.0x10^7$ $M^{-1}s^{-1}$ | | |
| 11 | VIIIa + IXa ↔ IXa = VIIIa | $1.0x10^7$ $M^{-1}s^{-1}$ | $5.0x10^{-3}$ $s^{-1}$ | |
| 12 | IXa = VIIIa + X ↔ IXa = VIIIa = X → IXa = VIIIa + Xa | $1.0x10^8$ $M^{-1}s^{-1}$ | $1.0x10^{-3}$ $s^{-1}$ | 8.2 $s^{-1}$ |
| 13 | VIIIa ↔ $VIIIa_1$ + $VIIIa_2$ | $6.0x10^{-3}$ $s^{-1}$ | $2.2x10^4$ $M^{-1}s^{-1}$ | |
| 14 | IXa = VIIIa = X → IXa + X + $VIIIa_1$ + $VIIIa_2$ | $1.0x10^{-3}$ $s^{-1}$ | | |
| 15 | IXa = VIIIa → IXa + $VIIIa_1$ + $VIIIa_2$ | $1.0x10^{-3}$ $s^{-1}$ | | |
| 16 | IIa + V → IIa + Va | $2.0x10^7$ $M^{-1}s^{-1}$ | | |
| 17 | Xa + Va ↔ Xa = Va | $4.0x10^8$ $M^{-1}s^{-1}$ | 0.2 $s^{-1}$ | |
| 18 | Xa = Va + II ↔ Xa = Va = II → Xa = Va + mIIa | $1.0x10^8$ $M^{-1}s^{-1}$ | 103 $s^{-1}$ | 63.5 $s^{-1}$ |
| 19 | Xa = Va + mIIa → Xa = Va + IIa | $1.5x10^7$ $M^{-1}s^{-1}$ | | |
| 20 | Xa + TFPI ↔ Xa = TFPI | $9.0x10^5$ $M^{-1}s^{-1}$ | $3.6x10^{-4}$ $s^{-1}$ | |
| 21 | TF = VIIa = Xa + TFPI ↔ TF = VIIa = Xa = TFPI | $3.2x10^8$ $M^{-1}s^{-1}$ | $1.1x10^{-4}$ $s^{-1}$ | |
| 22 | TF = VIIa + Xa = TFPI → TF = VIIa = Xa + TFPI | $5.0x10^7$ $M^{-1}s^{-1}$ | | |
| 23 | Xa + ATIII → Xa = ATIII | $1.5x10^3$ $M^{-1}s^{-1}$ | | |
| 24 | mIIa + ATIII → mIIa = ATIII | $7.1x10^3$ $M^{-1}s^{-1}$ | | |
| 25 | IXa + ATIII → IXa = ATIII | $4.9x10^2$ $M^{-1}s^{-1}$ | | |
| 26 | IIa + ATIII → IIa = ATIII | $7.1x10^3$ $M^{-1}s^{-1}$ | | |
| 27 | TF = VIIa + ATIII → TF = VIIa = ATIII | $2.3x10^2$ $M^{-1}s^{-1}$ | | |
| 28 | IXa + X → IXa + Xa | | $K_m$ = $1,4x10^{-7}$ M | $K_{cat}$ = 0,048 $s^{-1}$ |
